# The spatial coding of touch is defined in intrinsic, limb-specific coordinates: an EEG study

**DOI:** 10.1101/2025.06.03.657562

**Authors:** Valeria C. Peviani, Hüseyin O. Elmas, W. Pieter Medendorp, Luke E. Miller

## Abstract

The brain localizes touch in space by integrating tactile and proprioceptive signals, a process known as tactile remapping. While it is often assumed that the remapped touch is encoded in an extrinsic, limb-independent reference frame, an alternative view proposes that touch may instead be represented within an intrinsic, limb-specific coordinate system. To test these hypotheses, we used electroencephalography (EEG) and a novel tactile stimulation paradigm in which participants received touch on their hands positioned at various locations relative to the body. Previous findings suggest that neural activity in primate sensorimotor and parietal regions monotonically encodes limb position. We therefore analyzed amplitude gradients in somatosensory evoked potentials (SEPs) to test predictions from each coding scheme. If touch is coded extrinsically, neural gradients should reflect changes of the external stimulus location, regardless of the limb. If coded intrinsically, gradients should be tied to the position of each limb and mirror each other between hands. Both univariate and multivariate EEG analyses found no evidence for extrinsic coding. Instead, we observed neural signatures of limb-specific, intrinsic spatial codes, the earliest emerging about 160 ms after touch in centro-parietal regions, later shifting to fronto-temporal and parieto-occipital areas. Furthermore, a population-based neural network model of tactile remapping successfully reproduced the observed gradient patterns. These results show that the human brain localizes touch using an intrinsic, limb-specific spatial code, challenging the dominant assumption of extrinsic encoding in tactile remapping.

## Introduction

When a fly lands on our skin, we immediately perceive which body part it is touching (e.g., on the hand dorsum). However, if we want to effectively act on the fly—such as swiping it off— knowing which body part is not sufficient; instead, we need to perceive its location within the space around us (e.g., on our left side). To compute the fly’s location in space, the brain must combine tactile signals encoding its location on the body surface with proprioceptive signals encoding body posture. This process is known as *tactile remapping*, as the ‘somatotopic’ touch location on the skin surface is remapped into a ‘spatiotopic’ reference frame that specifies where the stimulus is in the space surrounding us (Medina and Coslett, 2010; Heed et al., 2015).

The crossed-hands paradigm is often used to investigate the neural underpinnings of tactile remapping. In this paradigm, tactile stimuli are typically delivered on the hands while they are either crossed or uncrossed over the body midline (e.g., Yamamoto and Kitazawa, 2001; Shore et al., 2002; Azañón and Soto-Faraco, 2008). This manipulation highlights the involvement of multimodal integration, as the somatotopic position of touch must be integrated with proprioceptive signals encoding arm posture to compute its spatiotopic position. Spatiotopic coding emerges within ∼100-150 ms after touch (Heed and Röder, 2010; Soto-Faraco and Azañón, 2013; Alouit et al., 2024) and involves processing in parietal and premotor cortices (e.g., Lloyd et al., 2003; Azañón et al., 2010; Buchholz et al., 2011; Crollen et al., 2017; Klautke et al., 2023; Fabio et al., 2024).

Despite the long history of research on tactile remapping, the organizational principles of spatiotopic reference frames (e.g., the coordinate geometry) have remained largely unexplored. Take the case where touch is applied to either the left or right hand, both of which could be positioned at various spatiotopic locations. One intuitive possibility is that touch on the hands is remapped to a *shared* spatiotopic reference frame (**Figure 1A, top**), centered on a common origin (e.g., the body midline, the gaze, Heed et al., 2015). In this view, the spatial position of touch is limb-independent—touch on the hand is encoded based on the hand position in physical space, regardless of which hand is touched. We will therefore refer to this type of remapping as *extrinsic coding*.

**Figure 1.**
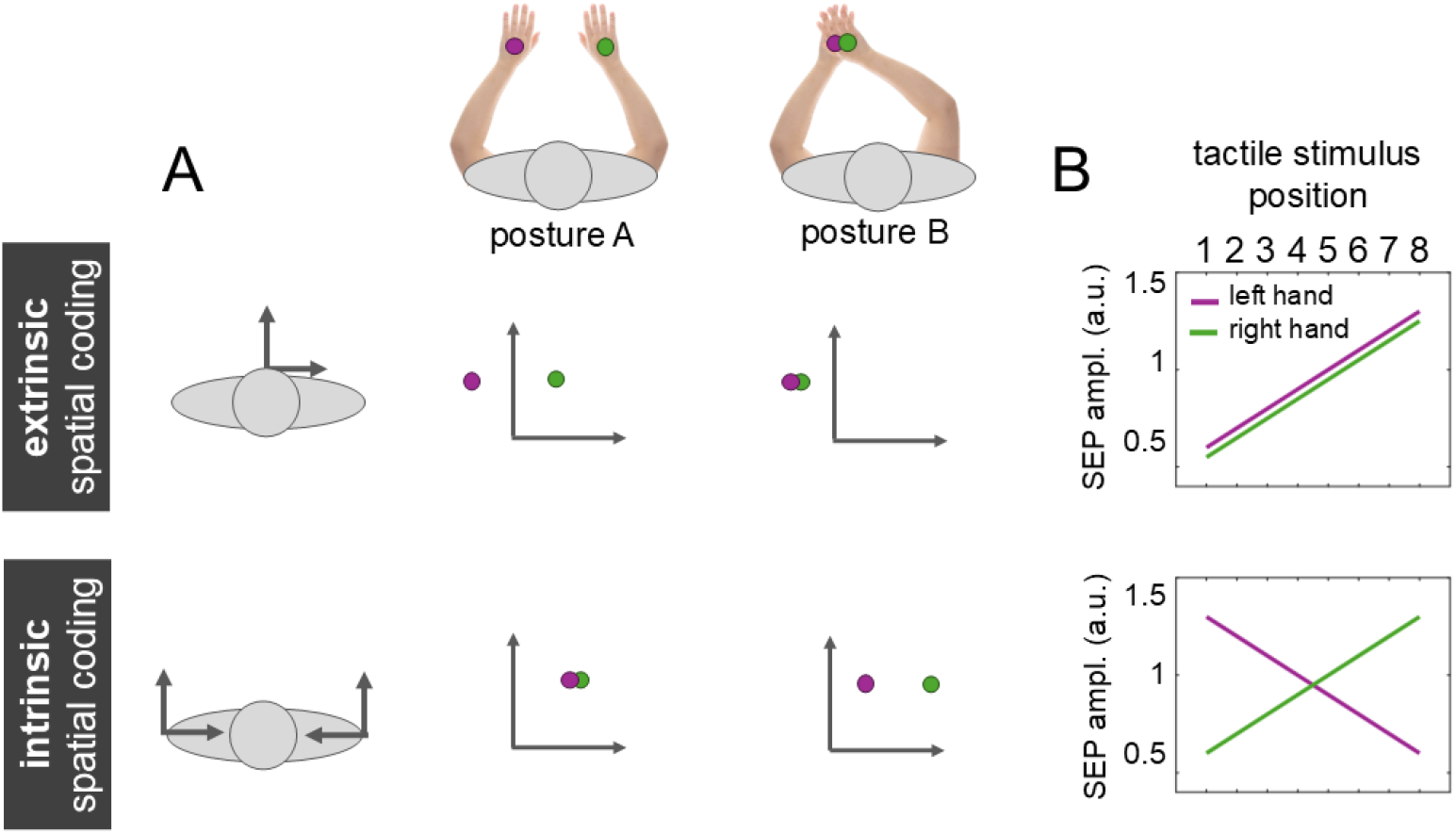
Extrinsic and intrinsic spatial coding. **A**. Examples of extrinsic (top row) and intrinsic (bottom row) spatial coding of tactile events (purple and red dots) occurring at different body postures. In the extrinsic coding, locations are defined based on a common origin, here placed at the body midline. Touch location is thus limb independent, as it is defined by its physical position relative to the origin, regardless of where it happens on the body. Tactile stimuli delivered on the hands are in the same extrinsic coordinates when the hands occupy the same position in physical space, like in Posture B. Instead, the intrinsic coding is tied to the joint kinematics of the limbs and is therefore defined by mirror-symmetric sets of coordinates, here centered at the shoulders. In this case (see Posture A), tactile stimuli delivered on the hands are in the same intrinsic coordinates when the arms are in the same posture (i.e., same patterns of joint angles). **B**. Predictions of SEP amplitude linear modulation as a function of tactile stimulus spatial location, with location 1 being at the level of the left shoulder, and 8 at the level of the right shoulder. For the extrinsic reference frame (top), the amplitude modulation is tied to the absolute position of the stimulus, regardless of which hand is touched. That is, the modulation gradient increases in the same direction for both hands (e.g., from position 1 to 8). For the intrinsic reference frame (bottom), the amplitude modulation is tied to each limb, assuming mirror symmetric properties. For instance, it may increase from the most contralateral position (1 for the right hand; 8 for the left hand) to the most ipsilateral (8 for the right hand; 1 for the left hand).

Alternatively, touch on each hand could be remapped into a limb-specific reference frame whose spatial coding is based on the current postural configuration of each limb (**Figure 1A, bottom**). That is, the organizational properties of the reference frame are tied to the biomechanical properties of each forelimb, such as the bilateral symmetry of limb geometry and joint flexion/extension. For this reason, we will refer to this type of remapping as *intrinsic coding*. Notably, this form of remapping uniquely predicts that the spatial coding of touch on the two limbs has *mirror symmetric* properties. This type of reference frame plays an important role in motor learning and reach control (Brayanov et al., 2012;).

In the present study, we used EEG to explore whether tactile remapping involves extrinsic and/or intrinsic spatial coding in the human brain. We designed a task in which touch was delivered to each hand, which occupied one of eight spatial locations relative to the body. Inspired by neurophysiological recordings in sensorimotor cortices (Lacquaniti et al., 1995; Tillery et al., 1996), we hypothesized that a monotonic spatial gradient in SEP amplitude reflects the spatiotopic coding of touch. Extrinsic coding predicts a spatial gradient tied to the absolute position of the stimulus, regardless of which hand is touched (**Figure 1B, top**).

Intrinsic coding instead predicts a mirror-symmetric gradient, tied to each hand independently (**Figure 1B, bottom**). Using both univariate and multivariate analyses, we observed the expected neural signature of intrinsic coding approximately 160 ms after touch. A population-based neural network based on properties of tactile and proprioceptive coding reproduced the observed mirror-symmetric spatial modulation. These findings suggest that tactile localization relies on intrinsic limb-specific spatiotopic representations rather than extrinsic limb-independent spatial coordinates.

## Materials and Methods

### Participants

Twenty-two healthy participants were recruited from the university pool to participate in the experiment; two were discarded due to technical issues (final N = 20, 19 females; age: 20.2 ± 2.4 years old). Sixteen participants were fully right-handed (laterality quotient: 100), two were ambidextrous (laterality quotients: 0 and 50) and one was left-handed (laterality quotient: -100), as assessed using the short version of the Edinburgh Handedness Scale (Oldfield, 1971; Veale, 2014). The study was approved by the ethics committee of the Faculty of Social Sciences at Radboud University (ECSW-2022-085R1). Participants gave their written informed consent before participating in the experiment.

### Task and experimental procedure

#### High density crossed hands paradigm

The participant sat at a table covered with a wooden panel with eight dots arranged along a horizontal line, each spaced 15 cm apart. The panel was centered on the participant’s midline. A computer screen was placed 50 cm in front of the participant and centered on their midline. At the beginning of each block, the participant was instructed to configure their left (LH) and right (RH) hand according to a visual representation of the eight locations displayed on the computer screen (**Figure 2A**).

**Figure 2.**
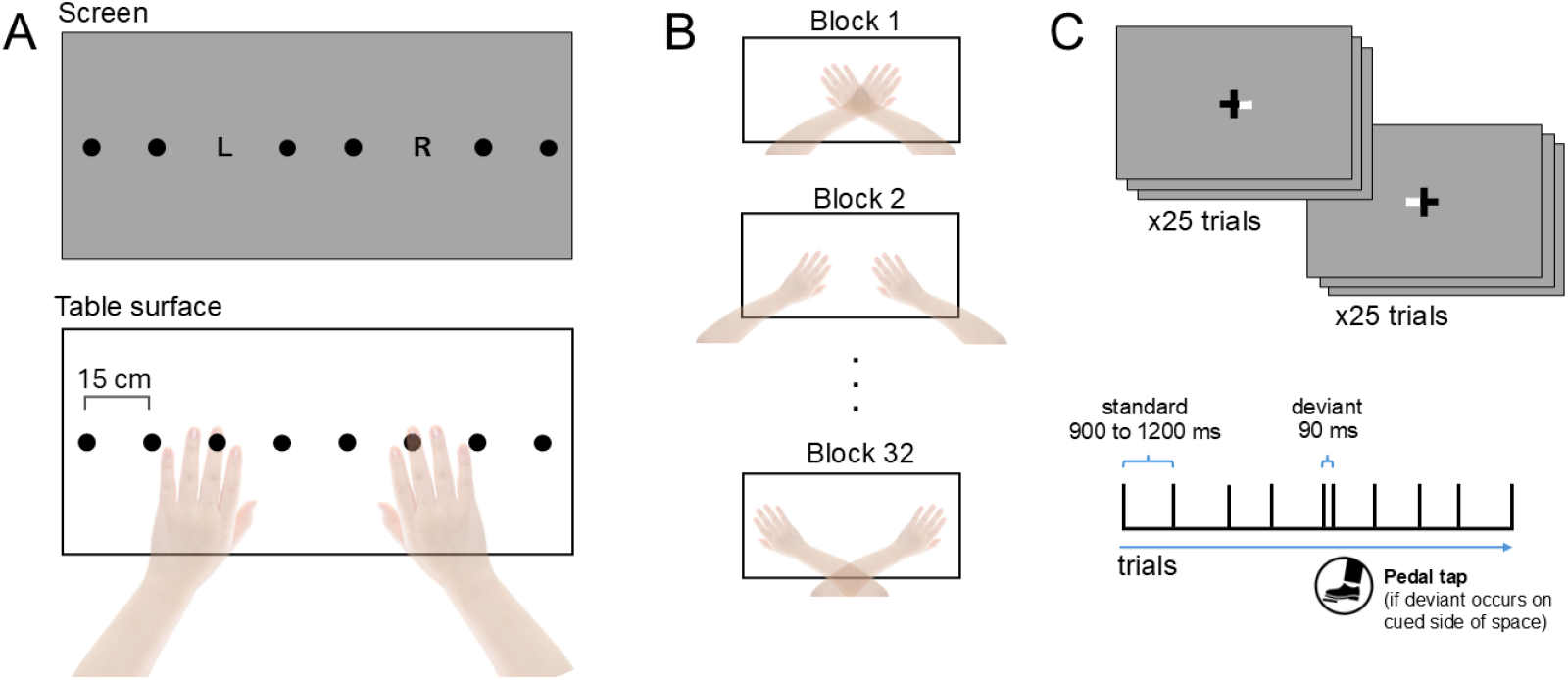
Experimental paradigm. **A**. At the beginning of each experimental block participants were instructed to position their hands on two out of eight locations marked on a horizontal panel, based on the instructions given on the screen. Specifically, the left and right middle fingers had to be placed at the positions taken by the L and R letters. **B**. The experiment included 32 blocks, each including 60 electrocutaneous stimulations and lasting around a minute each. **C**. In order to have participants paying attention to the stimulations, we used a somatosensory spatial attention task in which participants had to respond via pedal tap every time they detected a deviant stimulus (double stimulation) on the side of space cued by the fixation cross (white segment).

The LH and RH remained at the specified two of the eight locations throughout the duration of each experimental block. We tested eight different left-right hand configurations, all mirror-symmetric: i) LH in position 1 and RH in position 8; ii) LH in 2 and RH in 7; iii) LH in 3 and RH in 6; iv) LH in 4 and RH in 5; v) LH in 5 and RH in 4; vi) LH in 6 and RH in 3; vii) LH in 7 and RH in 2; viii) LH in 8 and RH in 1. Note that the arm postures are uncrossed for configurations i-iv and are crossed for configurations v-viii. Each configuration was repeated four times in a pseudo-random order for a total of 32 blocks (**Figure 2B**).

During each block, participants performed a somatosensory spatial attention task that required to discriminate standard and deviant electrocutaneous stimuli. This task was introduced to ensure that participants paid attention to the tactile stimulation. Stimuli were sequentially delivered to their left and right middle fingers, with an interstimulus interval randomly sampled from a uniform distribution with limits 900 to 1200 ms. The *standard* stimulus consisted of a single electrocutaneous stimulation (1 ms duration), while the *deviant* stimulus consisted of a double stimulation (i.e., two consecutive 1-ms stimulations spaced by 90 ms). Participants were instructed to fixate on a black cross positioned at the center of the computer screen (grey background). One of the horizontal segments of the fixation cross was white, indicating the side of space to direct their attention (**Figure 2C**). Participants were tasked with detecting deviant stimuli that occurred on the attended side of space and respond by tapping a foot pedal positioned beneath their right foot. The deviant trials were excluded from subsequent analysis.

Each block contained a total of 60 stimuli (30 delivered to the left hand and 30 to the right hand), with a 6:1 standard-to-deviant ratio in a pseudo-randomized order. The task included a total of 1920 stimuli, of which 1600 standard stimuli were included in the analysis. The side of the visual cue (left or right segment of the fixation cross) changed after every 25 stimuli. Every block lasted approximately one minute. Participants could take a break every 12 blocks. The task lasted approximately 40 minutes. Every participant completed two practice blocks before starting the task. Behavioral results show that the task effectively directed participants’ attention to the tactile stimuli. The median correct detection rate for target deviant stimuli (correctly detected/all deviant stimuli*100) was 82.2% (IQR 23.3).

### Electrocutaneous stimulation

The electrocutaneous stimuli were delivered to the left and right hands using a Digitimer 7A (Digitimer Ltd Hertfordshire, UK), controlled by a custom-made MATLAB script (R2024a, The MathWorks, Natick, Massachusetts). We used four ring electrodes, with the anodes placed on the proximal phalanges and the cathodes on the middle phalanges of the middle fingers. The amplitude for stimulation was determined for each individual participant based on their tactile threshold.

The threshold was computed using an ascending staircase procedure starting from 10 mA (200 V, 1 ms) to achieve six reversals (i.e., when participant felt the stimulation). After each reversal, stimulus amplitude was decreased by 50% (first two reversals), 25% (third and fourth reversal) or 12% (last two reversals) until participants could not feel the stimulation anymore. The amplitude was then increased again according to the same amplitude steps until the next reversal. Participants responded by button press (detected / non detected) using the hand that was not tested at that moment. The total threshold was calculated as the mean reversal value across hands, and increased by 50% (cf., Cardini et al., 2011; Nierhaus et al., 2015; Cardini and Longo, 2016; Christie et al., 2019) to compute the suprathreshold value used in the experiment. To ensure this level was ‘well above threshold’ (Holmes and Tamè, 2023), we verified that each suprathreshold stimulus evoked a clear perceptual sensation across ten consecutive trials. The amplitude range varied between 35 and 82 mA, with mean 56.3, std 11.8 mA.

### Electroencephalography procedure

#### EEG recordings

EEG data were recorded continuously (1000 Hz) using a 64 channel ActiCap system and Brainvision (Brain Products GmbH, Gilching, Germany). Impedance of all electrodes was kept at <20 kΩ. AFz served as ground and TP9 as online reference. Task control was performed using a MATLAB (R2024a) custom-script on the experimental control computer, which was synchronized and communicated with the EEG data recording system. The trial events were sent to the EEG data recording system via a parallel port.

#### Preprocessing of EEG signals

EEG signals were preprocessed using the Fieldtrip Toolbox (Oostenveld et al., 2011) and analyzed using MATLAB (R2022b). The preprocessing steps for each participant were as follows: the EEG signal was interpolated for noisy/missing channels using a spherical spline and high-pass filtered at 0.1 Hz. Artifacts due to electrocutaneous stimulation were removed by interpolating the 0 to 15 ms post-stimulus window with the average signal collected -25 to - 5 ms before and 20 to 40 ms after the stimulus. We performed ICA to reject eye-blinks and horizontal eye-movement components. We then epoched data into a time window of 800 ms, from 300 ms before to 500 ms after each electrocutaneous stimulation. We manually rejected trials that were contaminated by muscle artifacts or other forms of signal noise via visual inspection. We low-pass filtered the signals at 40 Hz, re-referenced the data to the average voltage across the scalp and performed baseline correction (subtraction method with baseline -200 to -1 ms before stimulus).

### Analyses

#### General Linear Modelling

We aimed to investigate whether coding of touch is organized in extrinsic or intrinsic coordinates, which we hypothesized would be reflected in spatial gradients of SEP responses. We therefore fit general linear models (GLMs) to identify whether SEP amplitudes were modulated by the spatial position of touch (1 to 8), in each type of coding (see above; **Figure 1**). Average SEPs were computed separately for each coordinate system (intrinsic, extrinsic), spatial position (1 to 8), channel, and participant, considering all standard trials.

Given that each coding type predicts specific patterns of spatial gradients for each hand (**Figure 1B**), we first fit linear models considering each hand individually. We binned the signal with a window of 20 ms and fitted a linear regression model for each participant, channel, hand, and time point with spatial position as predictor. The numerical value assigned to each position was anchored to the extrinsic spatial location (i.e., 1 is the most leftward and 8 is the most rightward positions). We included contralateral and midline channels in the analyses, for a total of 35 channels (28 contralateral, 7 midline channels). We compared the slope distributions to zero using one-sample t-tests and corrected for False Discovery Rate (FDR) over time and channels to control for Type-I error. Extrinsic coding predicts that the slopes of the spatial gradients for each hand will have a similar sign (e.g. positive), whereas intrinsic coding predicts opposite signed slopes for each hand’s spatial gradient (**Figure 1B**).

We next fit linear models average right and left hand SEPs, related to extrinsic and intrinsic reference frames. Collapsing across the hands allowed us to boost the signal-to-noise of the fits, providing a better characterization of the properties of the spatial gradients. To calculate the *extrinsic coordinates SEPs*, we grouped trials in which the left and right hands were stimulated at each extrinsic spatial coordinate. For instance, the signal obtained from left and right hand stimulation at position 2 were averaged together for each midline and contralateral channel (e.g., C2 and C1). To calculate *intrinsic coordinates SEPs*, we grouped trials in which the left or right hands were stimulated at each intrinsic coordinate. For instance, signal obtained from trials in which the left hand was stimulated in position 2 was averaged with the signal obtained from trials in which the right hand was stimulated at position 7, as these two extrinsic locations reflect similar intrinsic coordinates (**Figure 3**). For these models, the numerical coding of the stimulus location was treated as if touch was on the right hand (i.e., 1 is most contralateral, and 8 most ipsilateral).

**Figure 3.**
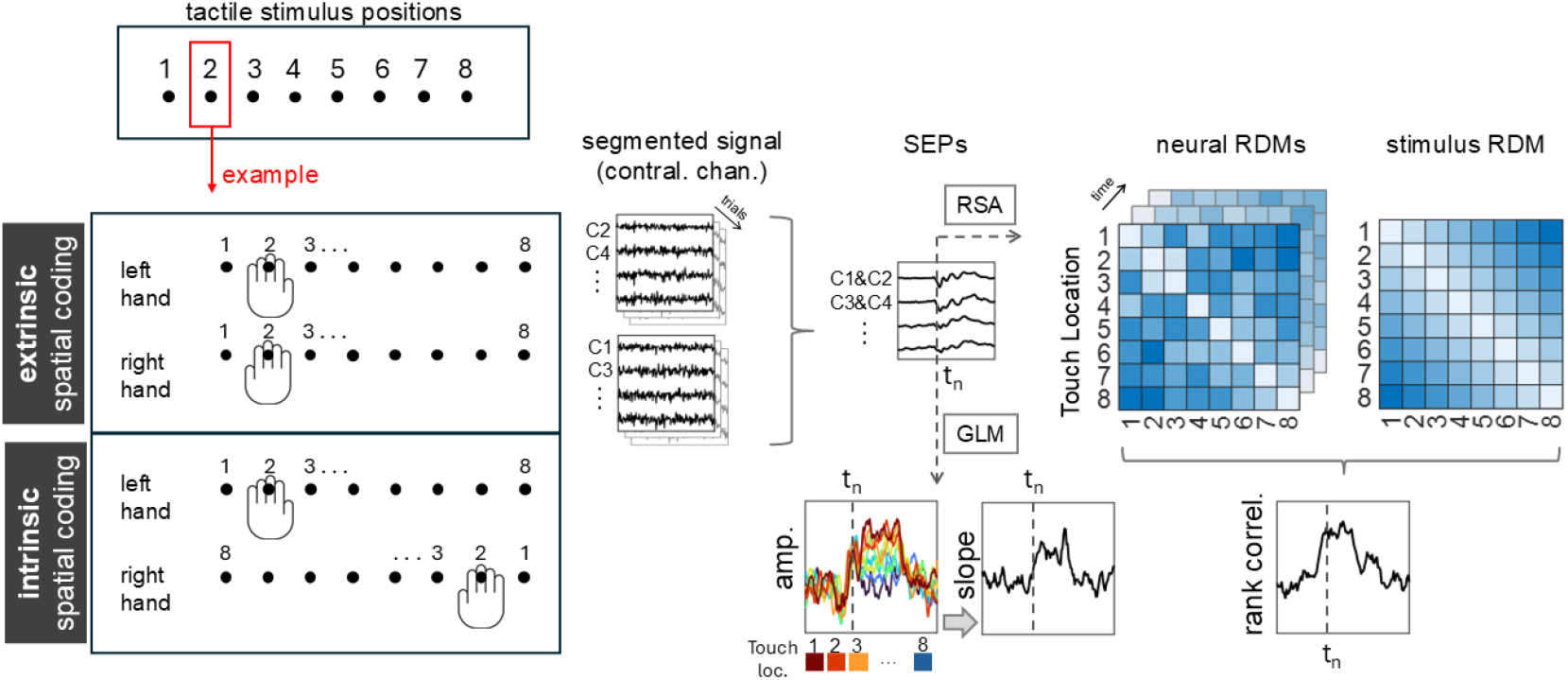
Analysis pipeline. EEG signal was preprocessed and segmented into trials starting 300 ms and ending 500 ms after the electrocutaneous stimulation onset. The analysis focused on channels contralateral to the stimulated hand. Trials were grouped based on the stimulus location considering two different reference frames (extrinsic, intrinsic). For the former, trials associated with left- and right-hand stimulations were grouped together when the hands were in the same extrinsic location (e.g., second location from the right). For the latter, trials were grouped together when the hands were in the same intrinsic location (e.g., left hand in the second location from the right, right hand in the second location from the left). Grouped trials were used to compute SEPs. We used general linear models (GLM) on each participant’s data to test for the effect of stimulus location (1 to 8) in each reference frame on the SEP amplitude at each time-point. We thus obtained a slope time-series for each channel, reference frame, and participant. Slopes were tested against zero and FDR correction was applied. We used representational similarity analysis (RSA) to correlate representational dissimilarity matrices (RDMs) obtained from the SEPs pertaining to each reference frame and the stimulus RDM specifying the distances between locations. Note that, while the neural RDM differs across reference frames, the stimulus RDM remains the same.

Even though we predicted a linear gradient given the neurophysiological findings (Kettner et al., 1988; Lacquaniti et al., 1995; Tillery et al., 1996), we also performed this analysis with several non-linear models as well (e.g. second-degree polynomials). However, these failed to perform better than the linear model. We therefore only report the results of our linear analyses.

#### Representational Similarity Analysis

We further analyzed the EEG data using Representational Similarity Analysis (RSA, Kriegeskorte et al., 2008) to detect similarities between the spatial configuration of touch locations and the patterns of electrophysiological activity. The stimulus representational dissimilarity matrix (RDM) was computed by taking the 1-dimensional Euclidean distances between each pair of touch locations. The neural RDMs were computed by taking the Euclidean distance (35-dimensional; i.e., the number of included channels) between the SEP amplitude obtained from each pair of touch location, for each participant, time point and spatial coding type (intrinsic, extrinsic). Note that the stimulus RDM was the same across reference frames because the physical distances between each pair of stimulus locations do not change. What allowed us to disentangle the contribution of intrinsic and extrinsic coordinate systems is how the neural RDM is calculated. Specifically, neural RDMs were based on averaged neural responses to left- and right-hand touches that share the same intrinsic or extrinsic coordinates. As a result, these neural RDMs differ depending on the reference frame being considered. See **Figure 3**. Stimulus and neural RDMs were correlated using Spearman’s rank correlation. The correlation time-series was binned with a window of 20 ms. For each time point, we used one-tailed Wilkinson signed-rank tests to compare the correlation values of each coordinate system with the noise floor, which leads to a more precise estimate of representational similarity given the noise in the data (Nili et al., 2014). The noise floor and ceiling represent the lower and upper edges of the average correlation of the stimulus RDM with the single-subject neural RDMs. The upper bound is estimated by computing the correlation between each subject’s RDM and the group-average RDM. The lower bound is estimated using a leave-one-out approach, where each participant’s RDM is correlated with the average RDM of the other participants’ RDMs. FDR correction was applied to control for Type-I error. We calculated the average variance explained by brain responses as: (correlation – lower noise bound)/(upper noise bound – lower noise bound).

To explore the topography of the neural and stimulus RDMs, we performed an additional RSA considering clusters of channels. In detail, we ran the RSA pipeline for each of the 35 channels, with the neural dissimilarity matrices calculated including 5 to 10 neighboring channels, defined using a Fieldtrip standard template. This provided a correlation time-series for each channel.

#### Neural network model

Tactile remapping arises from the combination of proprioceptive and tactile signals in parietal regions (e.g., Azanon et al., 2010; Heed et al., 2015). One influential proposal is that this integration operates through a multiplicative mechanism, similar to gain field transformations described in studies of coordinate remapping (e.g., Andersen et al., 1985; Salinas and Abbott, 1996; Pouget and Sejnowski, 1997; Avillac et al., 2005). Supporting this idea, previous work has shown multiplicative somatosensory responses in neural populations (Kim et al., 2015). To test the plausibility of this proposal in the context of our findings, we implemented a simple population-based neural network model designed to reproduce the observed spatial gradients in neural responses through tactile-proprioceptive integration.

We designed a two-layer neural network model that was inspired by previous work on gain fields and coordinate transformations (Salinas and Abbott, 1996). The first layer of the network contained two nodes/subpopulations: (i) a tactile node, based on the summed population activity of tactile neurons with bell-shaped tuning curves tuned to skin space; (ii) a proprioceptive node, whose tuning was tied to the hand’s lateral position within the workspace relative to the shoulder; crucially, the activity of this node was characterized by a monotonic linear spatial gradient in intrinsic coordinates. The second layer contained a single node whose activity was based on the product of the nodes of the previous layer. The activity of each node was stochastic, with additive Poisson noise (Fano factor of one).

For simplicity, the operations of the neural network were implemented at the population-level, focusing exclusively on the overall population firing rate. We simulated several hand positions and proprioceptive gradients.

## Results

Electrocutaneous stimulation leads to several well-defined SEPs in the first 500 ms after touch. Consistent with previous studies, we observed the expected early (N20, P45, N70), middle (P100, N140), and late-latency (P200, P300) components. Each component had the expected time course and scalp topography (**Figure 4**, Allison et al., 1991; Allison et al., 1992).

**Figure 4.**
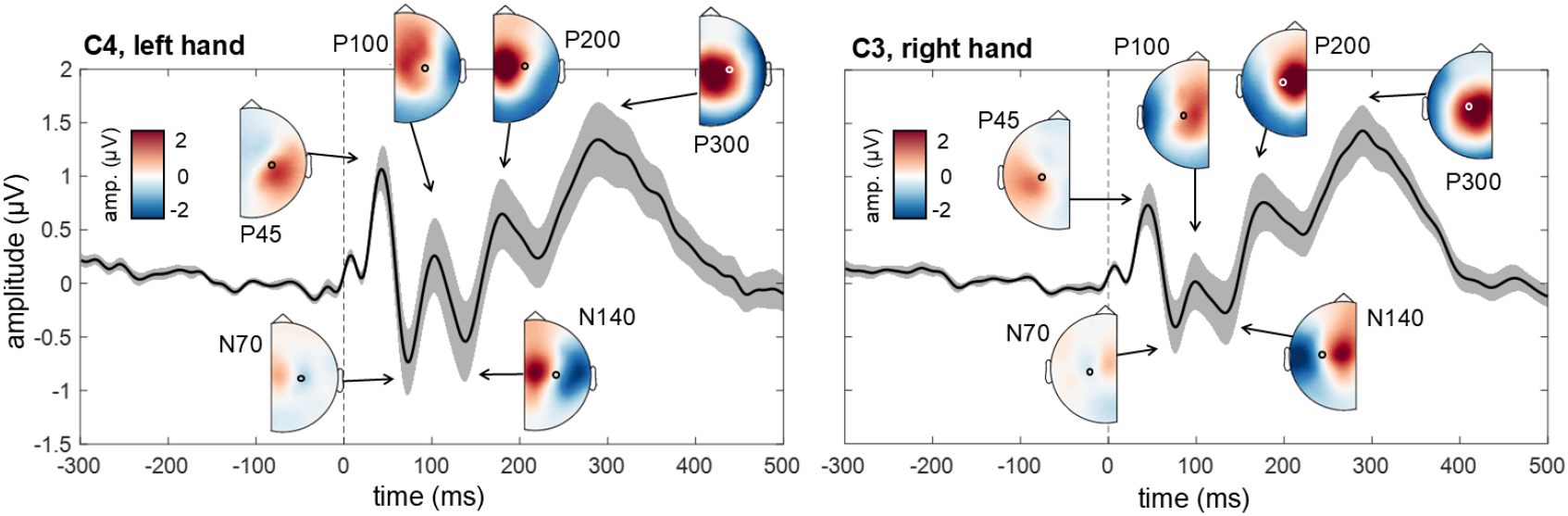
SEPs. Average SEPs obtained from left and right hand stimulations are plotted for a representative contralateral channel (C4 for the left hand, C3 for the right hand). Shaded area represents SEM. Scalp topography plots are shown for the main components.

We first used GLMs to characterize the reference frame during tactile localization. To do so, we investigated whether SEP amplitude was modulated by the spatial position of touch in either extrinsic or intrinsic coordinates. To test our specific predictions (**Figure 1B**), we first fit linear models to SEPs for each hand separately. Consistent with these predictions, we observed significant spatial gradients with opposite signs for each hand (**Figure 5**). Specifically, significant spatial gradients were observed within two different time-windows: from ∼160 to 300 ms mainly over central and centro-parietal channels; and from ∼320 to 460 ms over fronto-temporal, parietal and parieto-occipital channels. In neither instance did we find slopes that were significantly different from zero and had the same sign for both hands.

**Figure 5.**
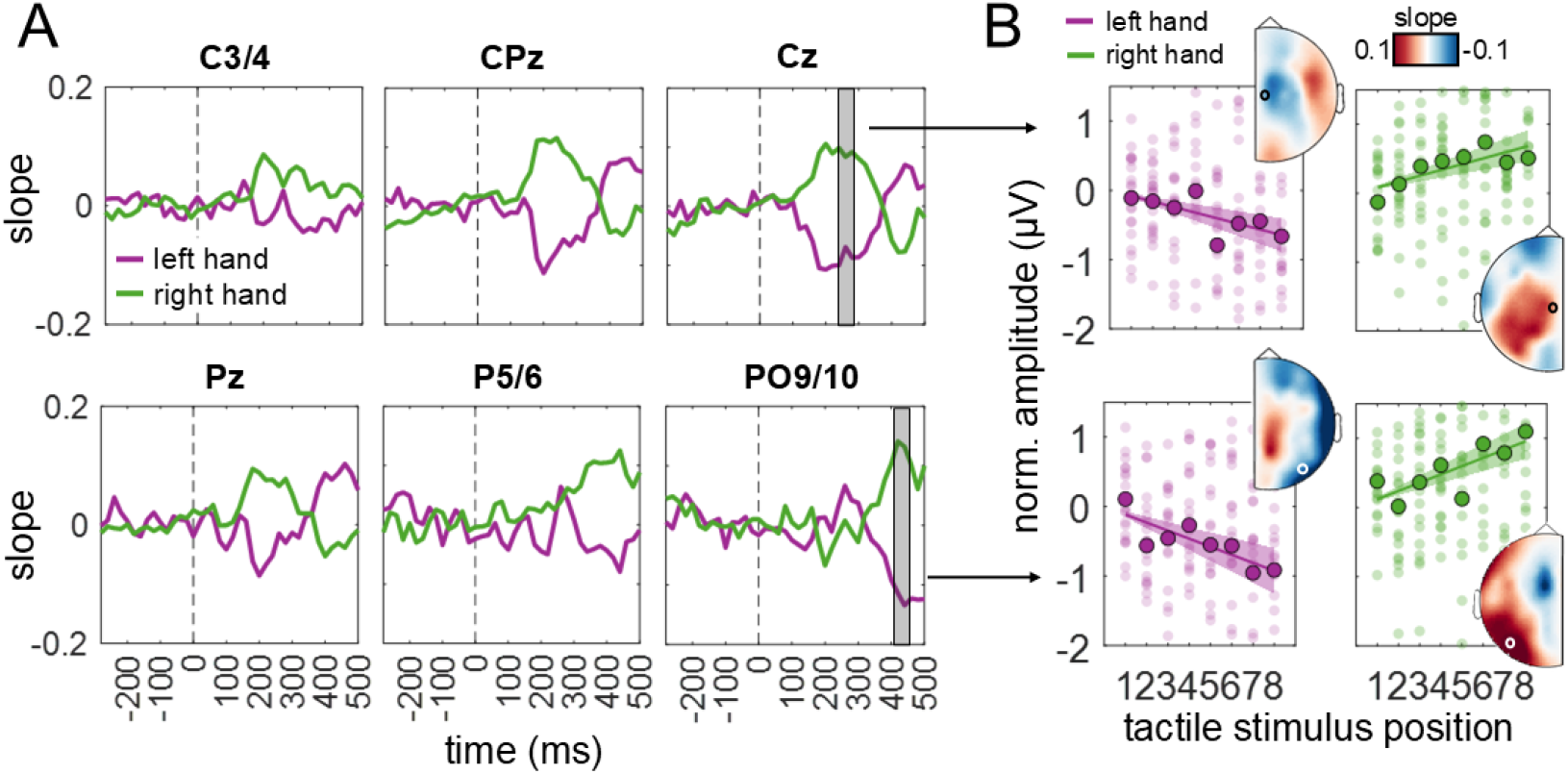
Mirror-symmetric spatial gradients for left and right hands. **A**. Slope time-series for left and right hands are plotted for six representative channels. Slopes were obtained from GLM fitted on each individual hand with stimulus position predicting SEP amplitude. **B**. Individual amplitude values (small dots) and mean amplitude values (big dots) are plotted against stimulus position. Note that amplitude values are normalized, i.e., the intercepts estimated through regression models were subtracted from the amplitude values at each time point. The regression lines (with SEM) illustrate the change in amplitude across positions for the corresponding time-windows marked in panel A. The amplitude from position 1 to 8 decreased for the left hand, while it increased for the right hand. In other words, the amplitude decreased for the most ipsilateral to the most contralateral positions.

To further characterize the spatial gradients, we next fit GLMs to SEP amplitudes averaged across hands in both intrinsic and extrinsic reference frames. As above, the location of touch had a significant relationship with SEP amplitude—but only when touch was defined within an intrinsic coordinate system. We again observed a significant spatial gradient within two different time-windows: from ∼160 to 300 ms mainly over central and centro-parietal channels; and from ∼320 to 460 ms over fronto-temporal, parietal and parieto-occipital channels. See **Figure 6A** for the slope time-series for two representative channels (C1/2 and P5/6) in the intrinsic coordinate system. For the extrinsic coordinate system, we found no modulation at any time-point in any channel on the scalp (**Figure 6B**).

**Figure 6.**
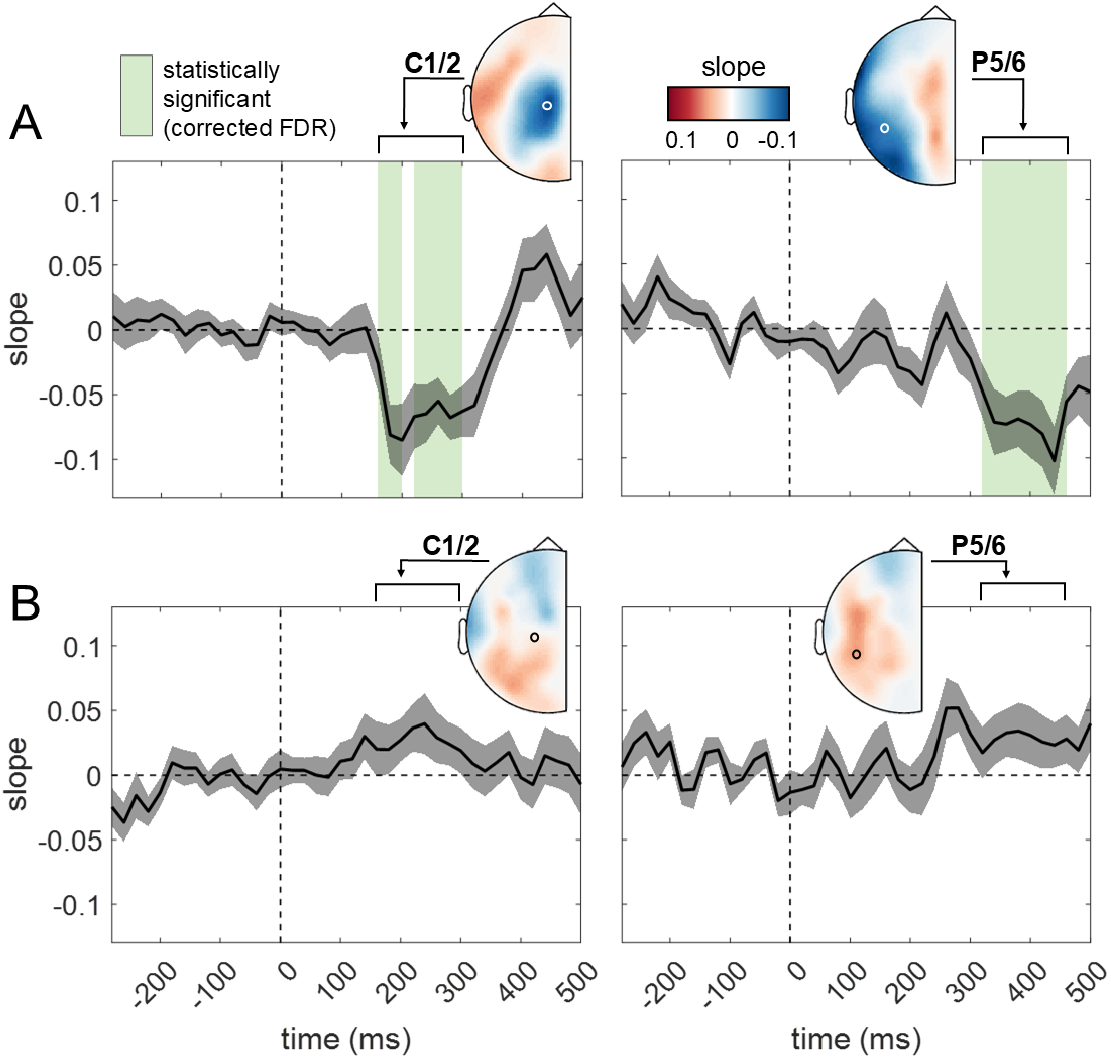
Slope time-series for the intrinsic (A) and extrinsic (B) coordinate systems. Slope time series are plotted for two representative channels, showing modulation based on stimulus spatial position encoded in intrinsic (A) or extrinsic (B) coordinates. The solid black line represents the group-level average, with the shaded gray area indicating SEM. Green vertical bands mark time windows where slope values significantly differ from zero (corrected for multiple comparisons using FDR), indicating significant amplitude modulation for the intrinsic coding. This modulation occurs from approximately 160 to 300 ms over centro-parietal channels (left plot) and later, from 320 to 460 ms, over parietal channels (right plot). Scalp topography maps display slope values over contralateral channels, averaged within these respective time windows, as indicated by the arrows.

This pattern of results reflects a modulation where SEP amplitude decreased as the hands moved from an uncrossed to a crossed posture. That is, neural responses were highest when the hands were directly in front of the shoulder (e.g., left hand in position 1) and linearly decreased as the hand was placed in the direction of the opposite shoulder (e.g., left hand in position 8). The observed spatial gradients were all negative, with an increase in amplitude up to 0.006 μV per cm. See **Figure 7** for the ERP amplitude modulation for C1/2 between 160 and 300 ms and P5/6 between 320 and 460 ms.

**Figure 7.**
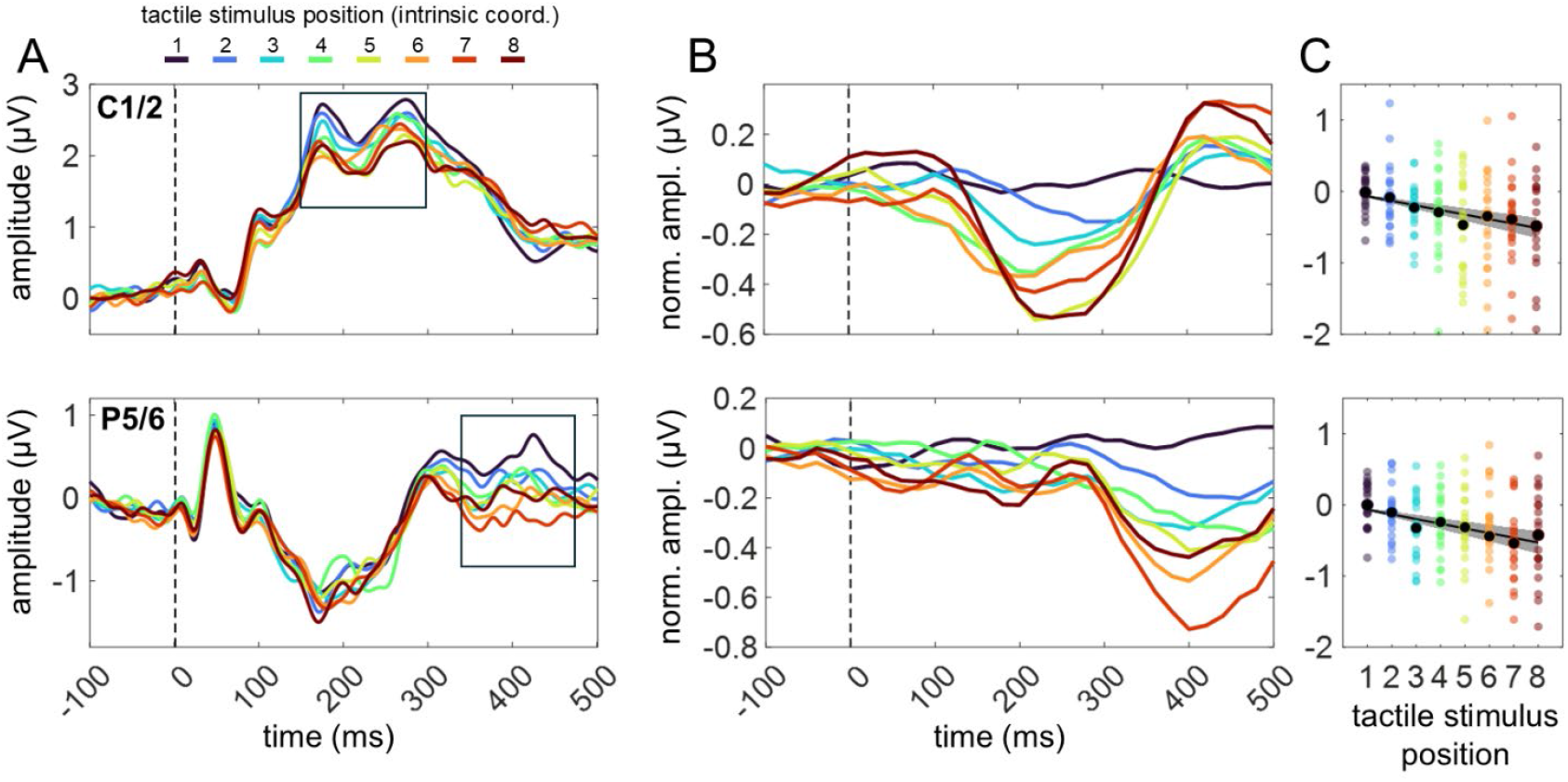
Amplitude modulation of stimulus position in intrinsic coordinates. **A**. Group-level SEPs for each stimulus position, encoded in intrinsic coordinates, are plotted for two representative channels (C1/2 top row, P5/6 bottom row). Different colors indicate different stimulus positions. Black boxes highlight time windows where regression analysis detected significant amplitude modulation, revealing a negative gradient in amplitude from position 1 (most leftward for the left hand, most rightward for the right hand) to position 8. **B**. To better visualize the gradient, the same signals are plotted after binning (20 ms window) and normalization, i.e., the intercepts estimated through regression models were subtracted from the amplitude values at each time point. **C**. Normalized individual amplitude values (colored dots) and mean amplitude values (black dots) and are plotted against intrinsic stimulus position. The regression line (with SEM) illustrates the decrease in amplitude across positions for the corresponding time-windows marked in panel A (160-300 ms for C1/2 and 320-460 ms for P5/6).

We next used RSA to identify spatial representations (extrinsic and/or intrinsic coding) in the multivariate patterns of somatosensory brain responses. In line with the above GLM analysis, we found evidence for the encoding of touch location in an intrinsic coordinate system, as indicated by a significant increase in ranked correlation relative to the noise floor between ∼180 and 400 ms after touch (**Figure 8A**). The average ± SEM variance explained by brain responses within such time window was 37.9 ± 10.5% (max 56.4%, ∼200 ms). The RSA performed on channel clusters provided a correlation time-series for each channel which we could show as a topographic map (**Figure 8A, scalp topographies**). Consistent with the GLM analysis, increased correlation values were initially observed in centro-parietal channels, before shifting towards fronto-temporal and parieto-occipital channels later in time. As before, we did not observe significant evidence for spatial coding in an extrinsic coordinate system (**Figure 8B**).

**Figure 8.**
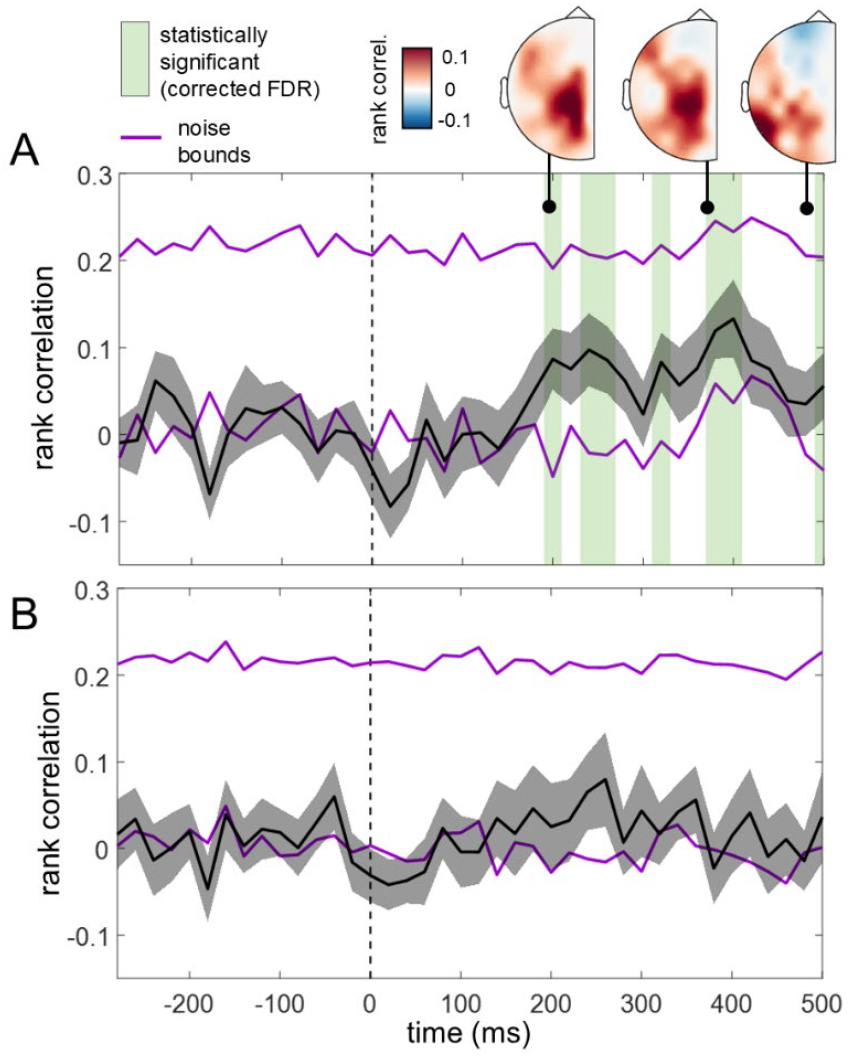
RSA results for the intrinsic (A) and extrinsic (B) coordinate system. The ranked correlation time series, averaged across participants, is shown in black (shaded area represents SEM). Purple lines represent noise bounds. Green vertical bands highlight time-windows where the rank correlation values were significantly greater than the lower noise bound (corrected FDR). In panel A, Scalp topography maps, obtained from an additional RSA based on channel clusters, illustrate the spatial distribution of rank correlation values at three selected time-points.

## Discussion

In this study, we combined a novel high-density crossed hands paradigm with simultaneous EEG recordings to investigate the organizational principles of the brain’s spatiotopic coding of touch. To do so, we measured somatosensory evoked brain responses to tactile stimuli presented on both hands, while the hands were placed at eight different spatial locations. We considered two possible organizational schemes for the spatial coding of touch (**Figure 1**): extrinsic coding, where the location of touch is encoded in a shared coordinate system that is independent of the limb; and intrinsic coding, tied to the mirror-symmetric biomechanics and proprioceptive coding of the limbs.

It is often assumed in the tactile remapping literature that the spatial coding of touch on the hands relies on a shared extrinsic spatial representation that is limb-independent (Heed et al., 2015). Our findings challenge this prevailing view. Both univariate and multivariate analyses provided compelling evidence that the spatial position of touch is encoded in intrinsic, rather than extrinsic coordinates. Using a GLM approach, we found that touch location is represented by an amplitude gradient in an intrinsic reference frame. Furthermore, representational similarity analysis (RSA) revealed a robust statistical relationship between spatial distances of touch locations in intrinsic space and patterns of neural activity. Thus, the location of touch is encoded in a limb-specific, intrinsic reference frame.

We identified significant intrinsic spatial coding starting around 160 ms after touch, a finding that is consistent with EEG studies using the crossed hands paradigm (Heed and Röder, 2010; Rigato et al., 2013; Soto-Faraco and Azañón, 2013; Alouit et al., 2024). The N140 and P200 are the most common SEP components exhibiting a postural modulation in their amplitudes. In keeping with other neurophysiological findings, these SEPs have sources within parietal and pre-central cortices (Arezzo et al., 1981; Allison et al., 1992; Desmedt and Tomberg, 1994; Hari and Forss, 1999; Waberski et al., 2002; Eimer and Forster, 2003). Our results suggest that these brain regions use an intrinsic spatial code for touch on the hands.

The SEP analysis identified a clear spatial gradient over the centro-parietal (∼160-300 ms), fronto-temporal and parieto-occipital (∼320-460 ms) channels. The spatial gradients were negative, meaning that SEP amplitude decreased as the hands moved from an uncrossed to a crossed posture. Furthermore, the neural responses were best fit with a linear model, indicating that SEP amplitudes monotonically decrease as the hand moved further away from the shoulder.

These results echo previous findings on the monotonic coding of hand location in single-unit recordings from monkeys, where neuronal firing rates in the motor cortex, somatosensory cortex, and posterior parietal cortex (Area 5) were found to vary linearly with the hand’s position in three-dimensional space (Kettner et al., 1988; Lacquaniti et al., 1995; Tillery et al., 1996). The parietal cortex, in particular, is thought to provide a spatial reference frame that supports the representation of space through position-dependent neural activity (Lacquaniti et al., 1995; Tillery et al., 1996; Medendorp and Heed, 2019). Our results extend these findings to tactile remapping by demonstrating that spatial gradients do not just encode hand position but are used to transform the somatotopic information into an intrinsic spatiotopic code. Furthermore, our analytic approach necessitates that the SEPs reflect non-linear integration between touch and hand position signals, as purely proprioceptive coding would have been removed by baseline correction. Thus, our results reflect the encoding of tactile space and not just hand position signals.

Along these lines, we propose that tactile remapping occurs via a multiplicative interaction between proprioceptive neurons and the skin-based neural processing in parietal regions. Multiplicative somatosensory responses have been observed previously (Kim et al., 2015), and are reminiscent of reference frame transformations involving gain fields (e.g., Andersen et al., 1985; Salinas and Abbott, 1996; Pouget and Sejnowski, 1997; Avillac et al., 2005). To demonstrate the plausibility of this proposal, we implemented a simple population-based neural network model that reproduced the observed spatial gradient in neural responses through tactile-proprioceptive integration (**Figure 9**). The tactile proprioceptive integration occurs via multiplicative interaction between bell-shaped cutaneous units and monotonically tuned proprioceptive units.

**Figure 9.**
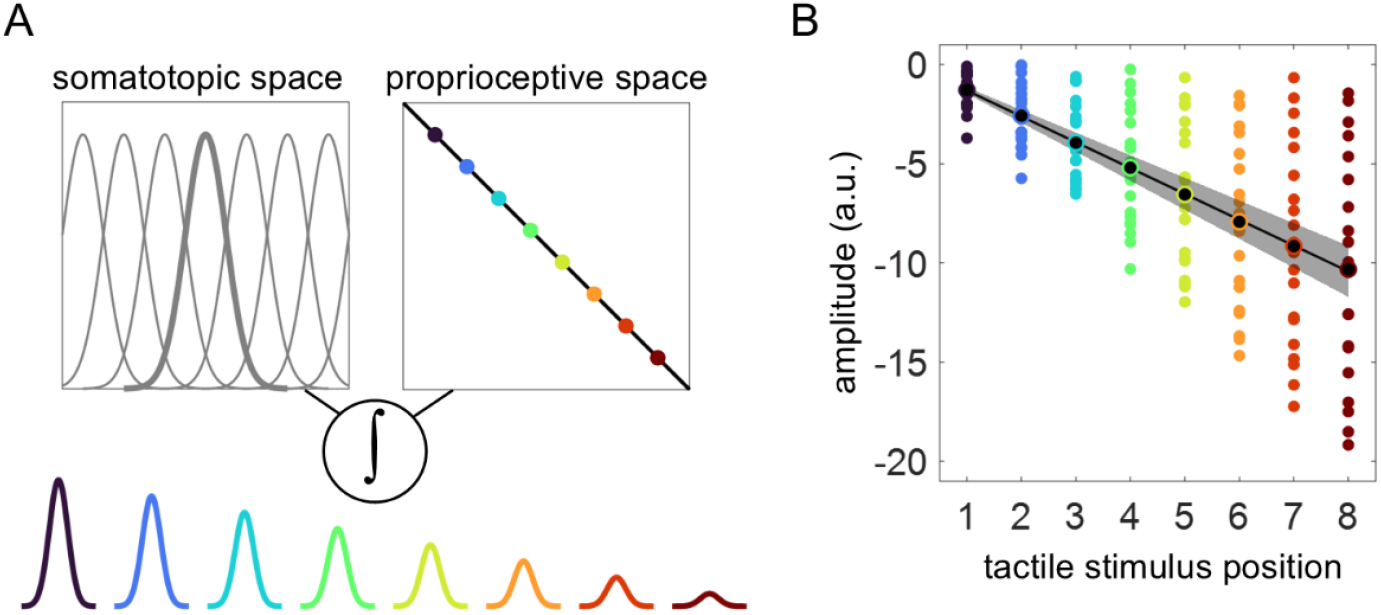
Population-based neural network for tactile-proprioceptive integration. **A**. The network includes two neural populations encoding touch and limb proprioception. The tactile unit encodes touch in a somatotopic space, (e.g., touch location on the hand surface) with uniform, bell-shaped firing rates across locations. The proprioceptive unit exhibits monotonic tuning of limb position, with firing rates varying proportionally with limb position (eight positions shown in different colors). The location of touch in space is specified by the product (∫) of the two population activities, leading to the response amplitude modulation. **B**. Simulated network responses for the eight tactile spatial locations used in the experiment. Each colored dot represents the average simulated response over 100 trials; black dots represent their grand means. The black regression line (with SEM) represents the average slope obtained from fitting GLMs with spatial locations predicting average responses.

Differences in SEP amplitudes between crossed and uncrossed postures have often been interpreted as evidence for a conflict between the somatotopic and spatiotopic reference frames (Rigato et al., 2013; Badde et al., 2014; Soto-Faraco and Azañón, 2013; Heed et al., 2015). This interpretation is based on idea that the crossed-hands posture (e.g., left hand on the right side of space) causes a misalignment between these two coordinate systems. Our findings suggest an alternative explanation: the SEP differences may simply reflect different spatial coordinates, encoded via the integration of tactile and proprioceptive signals (**Figure 9**). Consistent with previous studies (e.g., Soto-Faraco & Azanon, 2013), we showed that the amplitude difference along the tested posture is similar across the hands: Critically, this finding is compatible with an intrinsic, and not extrinsic, coding mechanism (**Figure 1**). This perspective moves beyond the idea of competing reference frames and instead emphasizes the dynamic interplay between somatosensory and postural cues.

Research on sensorimotor learning and control has shown that the motor system flexibly employs both intrinsic and extrinsic reference frames, switching between the two depending on task demands (Brayanov et al., 2012). For instance, proprioceptive-guided reaching seems to operate within intrinsic spatial maps (Sober and Sabes, 2003; Fujiwara et al., 2017). This may allow for an efficient mapping between proprioceptive signals and motor output. Coding touch on the forelimbs in intrinsic coordinates may similarly provide computational advantages. Consider again the example of using your left hand to swat off a fly on your right. If both the fly and the left hand were encoded in extrinsic coordinates, the reach would require a further transformation of the motor plan into intrinsic coordinates for movement control. However, coding the fly into intrinsic coordinates may circumvent the need for further remapping, reducing the computational steps for the sensorimotor control.

Nevertheless, a question that needs further investigation is if, and when, extrinsic coding emerges in tactile remapping. Motor control tasks that involve visuo-motor transformations are known to rely on extrinsic reference frames (Vetter et al., 1999; Krakauer et al., 2000; Ghez et al., 2007). Simple tasks that do not require visuomotor control, such as detecting deviant stimuli occurring on a specific side of space, may therefore only engage only the intrinsic reference frame. However, given the dynamic nature of spatial reference frames in the brain, it is plausible that variations of this task could elicit intrinsic-to-extrinsic coordinate transformation. For instance, if participants were asked to perform tasks requiring visuo-tactile integration or visuospatial transformations (e.g., localizing touch with visual pointer, matching touch location to a visual target), neural signatures of extrinsic coding might emerge. This possibility opens up promising avenues for future research.

Taken together, our findings provide theoretically significant insights into the organizational principles of spatiotopic coding in the human somatosensory system. We show that localization of touch relies on intrinsic, limb-specific reference frames, challenging the prevailing notion of limb-independent spatial coding. For the first time in humans, we describe positional gradients in SEPs over centro-parietal and fronto-temporal regions, which likely reflect mechanisms that have been only described in non-human primate studies. We extend these findings to tactile remapping, providing a plausible neural mechanism for encoding tactile space. Importantly, by showing progressive changes in amplitude as a function of touch spatial location, we move beyond the traditional binary comparison between postures (e.g., crossed-uncrossed) and stress the continuous integration between proprioceptive and tactile signals. In all, our study represents key groundwork for future investigations on tactile spatial maps in the brain.

## Contributions

VCP conceptualization, methodology, investigation, formal analysis, visualization, writing-original draft. HOE methodology, investigation. WPM conceptualization, methodology, writing-review and editing. LEM conceptualization, methodology, investigation, supervision, funding acquisition, writing-review and editing.

## Acknowledgements

LEM is supported by ERC 101076991 SOMATOGPS grant. WPM is supported by NWA-ORC-1292.19.298, NWO-SGW-406.21.GO.009 and Interreg NWE-RE:HOME. We are grateful to the members of the Brain, Body and Technology Lab for the insightful discussions.

## References

Allison T, McCarthy G, Wood CC (1992) The relationship between human long-latency somatosensory evoked potentials recorded from the cortical surface and from the scalp. Electroencephalography and Clinical Neurophysiology/Evoked Potentials Section 84:301– 314.

Allison T, McCarthy G, Wood CC, Jones SJ (1991) Potentials evoked in human and monkey cerebral cortex by stimulation of the median nerve: a review of scalp and intracranial recordings. Brain 114:2465–2503.

Alouit A, Gavaret M, Ramdani C, Lindberg PG, Dupin L (2024) Cortical activations associated with spatial remapping of finger touch using EEG. Cerebral Cortex 34:bhae161.

Andersen RA, Essick GK, Siegel RM (1985) Encoding of Spatial Location by Posterior Parietal Neurons. Science 230:456–458.

Arezzo JC, Vaughan HG, Legatt AD (1981) Topography and intracranial sources of somatosensory evoked potentials in the monkey. II. Cortical components. Electroencephalography and Clinical Neurophysiology 51:1–18.

Avillac M, Denève S, Olivier E, Pouget A, Duhamel J-R (2005) Reference frames for representing visual and tactile locations in parietal cortex. Nat Neurosci 8:941–949.

Azañón E, Longo MR, Soto-Faraco S, Haggard P (2010) The Posterior Parietal Cortex Remaps Touch into External Space. Current Biology 20:1304–1309.

Azañón E, Soto-Faraco S (2008) Changing Reference Frames during the Encoding of Tactile Events. Current Biology 18:1044–1049.

Badde S, Röder B, Heed T (2014) Multiple spatial representations determine touch localization on the fingers. Journal of Experimental Psychology: Human Perception and Performance 40:784–801.

Berniker M, Kording K (2008) Estimating the sources of motor errors for adaptation and generalization. Nat Neurosci 11:1454–1461.

Brayanov JB, Press DZ, Smith MA (2012) Motor Memory Is Encoded as a Gain-Field Combination of Intrinsic and Extrinsic Action Representations. J Neurosci 32:14951–14965.

Buchholz VN, Jensen O, Medendorp WP (2011) Multiple Reference Frames in Cortical Oscillatory Activity during Tactile Remapping for Saccades. J Neurosci 31:16864–16871.

Cardini F, Longo MR (2016) Congruency of body-related information induces somatosensory reorganization. Neuropsychologia 84:213–221.

Cardini F, Longo MR, Haggard P (2011) Vision of the Body Modulates Somatosensory Intracortical Inhibition. Cerebral Cortex 21:2014–2022.

Christie BP, Graczyk EL, Charkhkar H, Tyler DJ, Triolo RJ (2019) Visuotactile synchrony of stimulation-induced sensation and natural somatosensation. J Neural Eng 16:036025.

Crollen V, Lazzouni L, Rezk M, Bellemare A, Lepore F, Collignon O (2017) Visual Experience Shapes the Neural Networks Remapping Touch into External Space. J Neurosci 37:10097– 10103.

Desmedt JE, Tomberg C (1994) Transient phase-locking of 40 Hz electrical oscillations in prefrontal and parietal human cortex reflects the process of conscious somatic perception. Neuroscience Letters 168:126–129.

Eimer M, Forster B (2003) Modulations of early somatosensory ERP components by transient and sustained spatial attention. Exp Brain Res 151:24–31.

Fabio C, Salemme R, Farnè A, Miller LE (2024) Alpha oscillations reflect similar mapping mechanisms for localizing touch on hands and tools. iScience 27,3.

Fujiwara Y, Lee J, Ishikawa T, Kakei S, Izawa J (2017) Diverse coordinate frames on sensorimotor areas in visuomotor transformation. Sci Rep 7:14950.

Ghez C, Scheidt R, Heijink H (2007) Different Learned Coordinate Frames for Planning Trajectories and Final Positions in Reaching. Journal of Neurophysiology 98:3614–3626.

Hari R, Forss N (1999) Magnetoencephalography in the study of human somatosensory cortical processing. Philos Trans R Soc Lond B Biol Sci 354:1145–1154.

Heed T, Buchholz VN, Engel AK, Röder B (2015) Tactile remapping: from coordinate transformation to integration in sensorimotor processing. Trends in Cognitive Sciences 19:251–258.

Heed T, Röder B (2010) Common Anatomical and External Coding for Hands and Feet in Tactile Attention: Evidence from Event-related Potentials. Journal of Cognitive Neuroscience 22:184–202.

Holmes NP, Tamè L (2023) Detection, Discrimination & Localization: The Psychophysics of Touch. In: Somatosensory Research Methods (Holmes NP, ed), pp 3–33. New York, NY: Springer US.

Kettner RE, Schwartz AB, Georgopoulos AP (1988) Primate motor cortex and free arm movements to visual targets in three-dimensional space. III. Positional gradients and population coding of movement direction from various movement origins. J Neurosci 8:2938– 2947.

Kim SS, Gomez-Ramirez M, Thakur PH, Hsiao SS (2015) Multimodal Interactions between Proprioceptive and Cutaneous Signals in Primary Somatosensory Cortex. Neuron 86:555– 566.

Klautke J, Foster C, Medendorp WP, Heed T (2023) Dynamic spatial coding in parietal cortex mediates tactile-motor transformation. Nat Commun 14:4532.

Krakauer JW, Pine ZM, Ghilardi M-F, Ghez C (2000) Learning of Visuomotor Transformations for Vectorial Planning of Reaching Trajectories. J Neurosci 20:8916–8924.

Kriegeskorte N, Mur M, Bandettini PA (2008) Representational similarity analysis-connecting the branches of systems neuroscience. Front Syst Neurosci 2, 249.

Lacquaniti F, Guigon E, Bianchi L, Ferraina S, Caminiti R (1995) Representing Spatial Information for Limb Movement: Role of Area 5 in the Monkey. Cereb Cortex 5:391–409.

Lloyd DM, Shore DI, Spence C, Calvert GA (2003) Multisensory representation of limb position in human premotor cortex. Nat Neurosci 6:17–18.

Malfait N, Shiller DM, Ostry DJ (2002) Transfer of Motor Learning across Arm Configurations. J Neurosci 22:9656–9660.

Medendorp WP, Heed T (2019) State estimation in posterior parietal cortex: Distinct poles of environmental and bodily states. Progress in Neurobiology 183:101691.

Medina J, Coslett HB (2010) From maps to form to space: Touch and the body schema. Neuropsychologia 48:645–654.

Nierhaus T, Forschack N, Piper SK, Holtze S, Krause T, Taskin B, Long X, Stelzer J, Margulies DS, Steinbrink J, Villringer A (2015) Imperceptible Somatosensory Stimulation Alters Sensorimotor Background Rhythm and Connectivity. J Neurosci 35:5917–5925.

Nili H, Wingfield C, Walther A, Su L, Marslen-Wilson W, Kriegeskorte N (2014) A Toolbox for Representational Similarity Analysis. PLOS Computational Biology 10:e1003553.

Oldfield RC (1971) The assessment and analysis of handedness: The Edinburgh inventory. Neuropsychologia 9:97–113.

Oostenveld R, Fries P, Maris E, Schoffelen J-M (2011) FieldTrip: Open Source Software for Advanced Analysis of MEG, EEG, and Invasive Electrophysiological Data. Computational Intelligence and Neuroscience 2011:156869.

Pouget A, Sejnowski TJ (1997) Spatial Transformations in the Parietal Cortex Using Basis Functions. Journal of Cognitive Neuroscience 9:222–237.

Rigato S, Bremner AJ, Mason L, Pickering A, Davis R, van Velzen J (2013) The electrophysiological time course of somatosensory spatial remapping: vision of the hands modulates effects of posture on somatosensory evoked potentials. European Journal of Neuroscience 38:2884–2892.

Salinas E, Abbott LF (1996) A model of multiplicative neural responses in parietal cortex. Proc Natl Acad Sci USA 93:11956–11961.

Shadmehr R, Mussa-Ivaldi FA (1994) Adaptive representation of dynamics during learning of a motor task. J Neurosci 14:3208–3224.

Shore DI, Spry E, Spence C (2002) Confusing the mind by crossing the hands. Cognitive Brain Research 14:153–163.

Sober SJ, Sabes PN (2003) Multisensory Integration during Motor Planning. J Neurosci 23:6982–6992.

Soto-Faraco S, Azañón E (2013) Electrophysiological correlates of tactile remapping. Neuropsychologia 51:1584–1594.

Tillery SI, Soechting JF, Ebner TJ (1996) Somatosensory cortical activity in relation to arm posture: nonuniform spatial tuning. Journal of Neurophysiology 76:2423–2438.

Veale JF (2014) Edinburgh Handedness Inventory – Short Form: A revised version based on confirmatory factor analysis. Laterality 19:164–177.

Vetter P, Goodbody SJ, Wolpert DM (1999) Evidence for an Eye-Centered Spherical Representation of the Visuomotor Map. Journal of Neurophysiology 81:935–939.

Waberski TD, Gobbelé R, Darvas F, Schmitz S, Buchner H (2002) Spatiotemporal Imaging of Electrical Activity Related to Attention to Somatosensory Stimulation. NeuroImage 17:1347– 1357.

Yamamoto S, Kitazawa S (2001) Reversal of subjective temporal order due to arm crossing. Nat Neurosci 4:759–765.

